# An Optimal Policy Model for Concurrent Uncertainty Estimation During Decision Making

**DOI:** 10.1101/2021.10.14.464349

**Authors:** Xiaodong Li, Ruixin Su, Yilin Chen, Tianming Yang

## Abstract

We often postpone or even avoid making decisions when we feel uncertain. Uncertainty estimation is not an afterthought of decision making but a dynamic process that accompanies decision making in parallel and affects decision making. To study concurrent uncertainty estimation during decision making, we adapted the classic random-dots motion direction discrimination task to allow a reaction-time measure of uncertainty responses. Subjects were asked to judge whether a patch of random dots was moving left or right. In addition, they could seek assistance by choosing to look at a second stimulus that had the same direction but high coherence any time during the task. The task allows us to measure the reaction time of both the perceptual decisions and the uncertainty responses. The subjects were more likely to choose the uncertainty response when the motion coherence was low, while their reaction times of the uncertainty responses showed individual variations. To account for the subjects’ behavior, we created an optimal policy decision model in which decisions are based on the value functions computed from the accumulated evidence using a drift-diffusion process. Model simulations captured key features of the subjects’ choices, reaction times, and proportions of uncertainty responses. Varying model parameters explained individual variations in the subjects and the correlations between decision accuracy, proportions of uncertainty responses, and reaction times at the individual level. Our model links perceptual decisions and value-based decisions and indicates that concurrent uncertainty estimation may be based on comparisons between values of uncertainty responses and perceptual decisions, both of which may be derived from the same evidence accumulation process during decision making. It provides a theoretical framework for future investigations, including the ones that aim at the underlying neural mechanism.

## Introduction

Uncertainty estimation is part of decision making. For example, in a test in which cheat sheets are allowed, a student may directly answer a question if it seems easy. Alternatively, she may resort to the cheat sheet if she feels uncertain, sometimes well before she has formed any answers. The decision to look at the cheat sheet may be made immediately after she reads the question or after she spends a few minutes collecting her thoughts first. Whether and when to look at the cheat sheet is the result of a dynamic uncertainty estimation process that parallels and affects the decision-making process.

This concurrent uncertainty estimation during decision is distinct from most previous research studying uncertainty and confidence. Typically in these studies, the estimation of uncertainty or confidence is associated with a particular decision, and therefore made after a decision has been formed (Kepecs et al., 2008; Kiani et al., 2014; Kiani & Shadlen, 2009; Lak et al., 2014; Masset et al., 2020). For example, Kiani and Shadlen trained monkeys to perform a random dot motion direction discrimination task and offered them a third option (sure target), which provided a certain but smaller reward, to probe the monkeys’ confidence (Kiani & Shadlen, 2009). The monkeys chose the sure target more often when the motion stimulus was noisier. The sure target was only provided after the monkeys viewed the motion stimulus, thus the decision for the sure target was made based on an estimation of the accuracy of the perceptual decision. In another study, also based on the random dot motion discrimination task, human subjects were asked to point to horizontal bars to indicate both their choice and the decision certainty simultaneously (Kiani et al., 2014). Although the responses indicated both the perceptual decision and the decision certainty simultaneously, the certainty judgments were entangled with the perceptual decision and thus most likely made after the perceptual decisions. These established behavioral paradigms are inadequate to dissociate uncertainty estimation from decision outcomes, calling for new paradigms to investigate concurrent uncertainty estimation.

To this end, we adapted the classic random dot motion discrimination experiment as follows. When subjects were viewing the random dot motion and pondering upon its direction, in addition to making a direct perceptual decision, they had the option to seek assistance by asking for a second stimulus of the same direction but with a high motion coherence, which allowed a near-perfect decision accuracy. The decision for choosing the optional assistance is termed uncertainty response (UR), as the subjects should make such a decision only when they are uncertain about the first stimulus. The subjects were indeed found to make URs more often when the first stimulus had lower motion coherences. The reaction times of the UR, however, were not significantly affected by the motion coherence. In addition, there were significant variations in the reaction times among the subjects: some subjects’ reaction times of the UR were comparable to those of the perceptual decisions, while others’ reaction times were significantly longer.

To account for these results, we built an optimal policy decision model in which perceptual decisions and URs were both based on value functions computed from the accumulated evidence (Drugowitsch et al., 2012; Tajima et al., 2016, 2019). The model replicated the choice and reaction time patterns in both the perceptual decisions and the URs. Importantly, by varying model parameters, we were able to capture the behavior variability among the subjects and explain the interaction between the choice accuracies, UR proportions, and reaction times across the subjects. Our work establishes a reaction-time behavior paradigm for studying concurrent uncertainty estimation and provides a theoretical framework for both behavioral and future neurophysiology experiments aimed at studying the underlying neural mechanisms.

## Results

### Behavior task and Performance

Participants performed two versions of a random-dot motion direction discrimination task (**Figure 1**). In the standard task, the subjects were required to report the direction of a random dot motion stimulus (left vs. right) by pressing either the left or the right key whenever ready (**Figure 1A**). The strength of the motion was 0, 2, 4, 6, 8, 10, 12, 16, or 26% of coherence and varied randomly from trial to trial. In the second task, termed optional assistance task, when viewing the random dots, in addition to making a direct left or right decision (perceptual decision, PD), the subjects had the extra option of releasing and pressing the down key quickly again to get assistance from a second random dot stimulus (uncertainty response, UR). The second stimulus had the same motion direction as the first stimulus but a high coherence at 26% (**Figure 1B**). With the assistance from the second less noisy stimulus, the subjects, then, needed to report the direction of motion (perceptual decision in the second stage, PD2).

**Figure 1.**
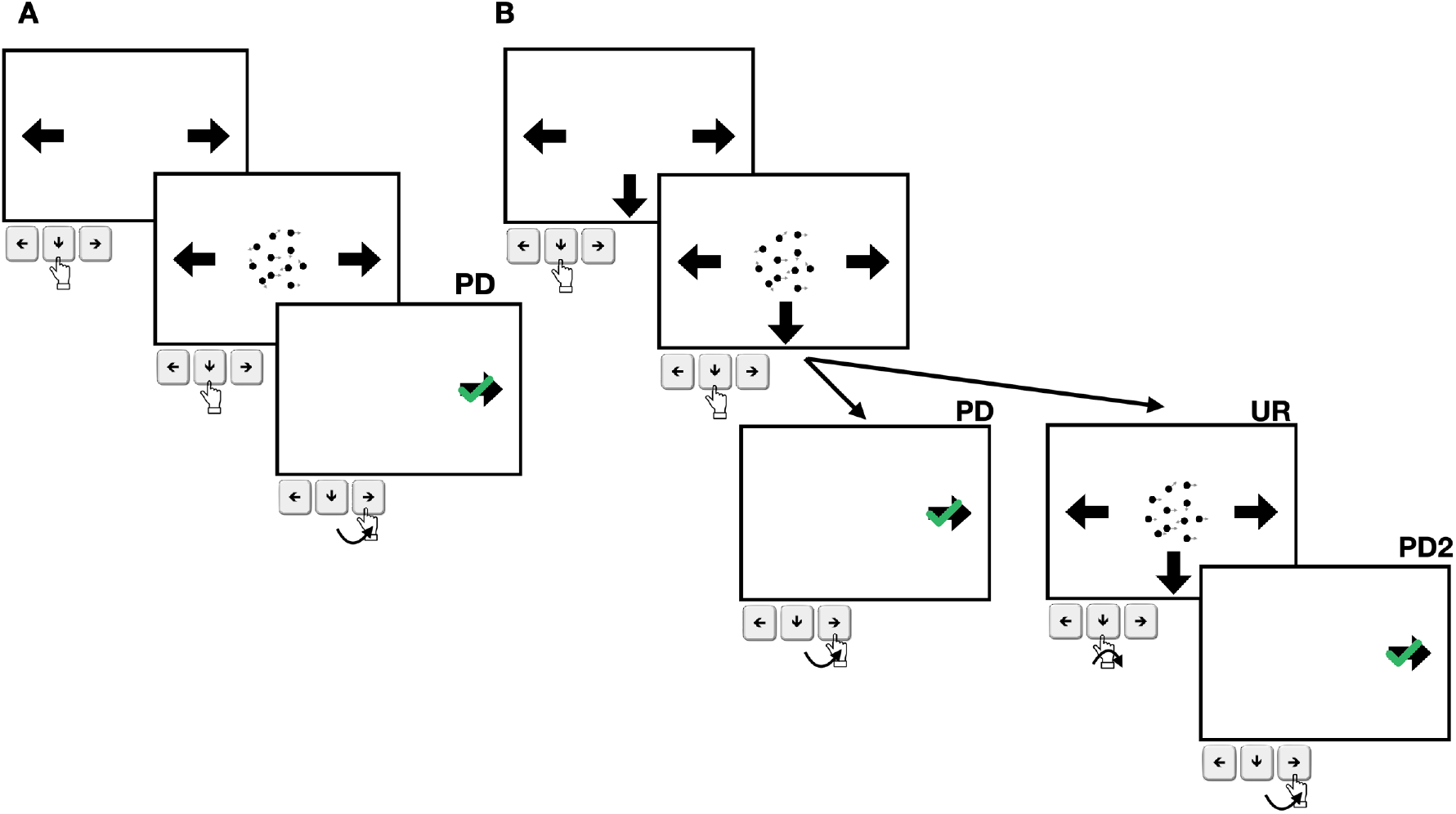
Task paradigms. **A**. Standard task. Two arrows appeared on the left and the right side of the screen to indicate the standard task. The subject pressed and held the down arrow key to start a trial. A patch of random dot motion stimulus was then presented at the center. The subject viewed the stimulus and released the down arrow key to press the left or the right arrow key to indicate her choice at any time. A green check indicated a correct response, and a red cross indicated an error. **B**. Optional assistance task. An additional down arrow was initially displayed on the screen to indicate the optional assistance task condition. When viewing the motion stimulus, in addition to pressing the left or right key to indicate her perceptual decision (PD) directly, the observer was allowed to release and press the down arrow key quickly again (within 200 ms) to view a second stimulus (UR). The observer, after viewing the second stimulus, had to release the down arrow key and press the left or the right arrow key to indicate the left or right choice (PD2).

In both tasks, the subjects performed better at higher motion coherences (**Figure 2A, Supplementary Figure S1** left column). When the assistance option was available but the subjects forwent the assistance and made a PD directly, their decisions were slightly, but significantly more accurate than their decisions in the standard task (Wilcoxon test, p<0.01). The subjects were near perfect in trials that they chose to look at the second stimulus (**Figure 2A, Supplementary FiguresS1**, left column). The PD reaction times (**Figure 2B, Supplementary Figure S1**, center column) in both tasks depended on the stimulus strength: weaker and more difficult stimuli led to longer reaction times. The overall reaction times of the PDs were longer in the standard task than in the optional assistance task. The reaction times of the UR, on average, were slower than those of the PDs in the optional assistance task. In contrast to the reaction times of the PDs, the reaction times of the UR did not appear to depend on the stimulus coherence, neither in the subjects’ average (**Figure 2B**, H_0_: slope=0, p=0.21) or in any individual subjects (**Supplementary Figure S2**). The PD2’s reaction times did not correlate with the first stimulus’s coherence (**Figure 2D**). Finally, as expected, the subjects were more likely to seek assistance when the motion coherence was lower (**Figure 2C, Supplementary Figure S1**, right column).

**Figure 2.**
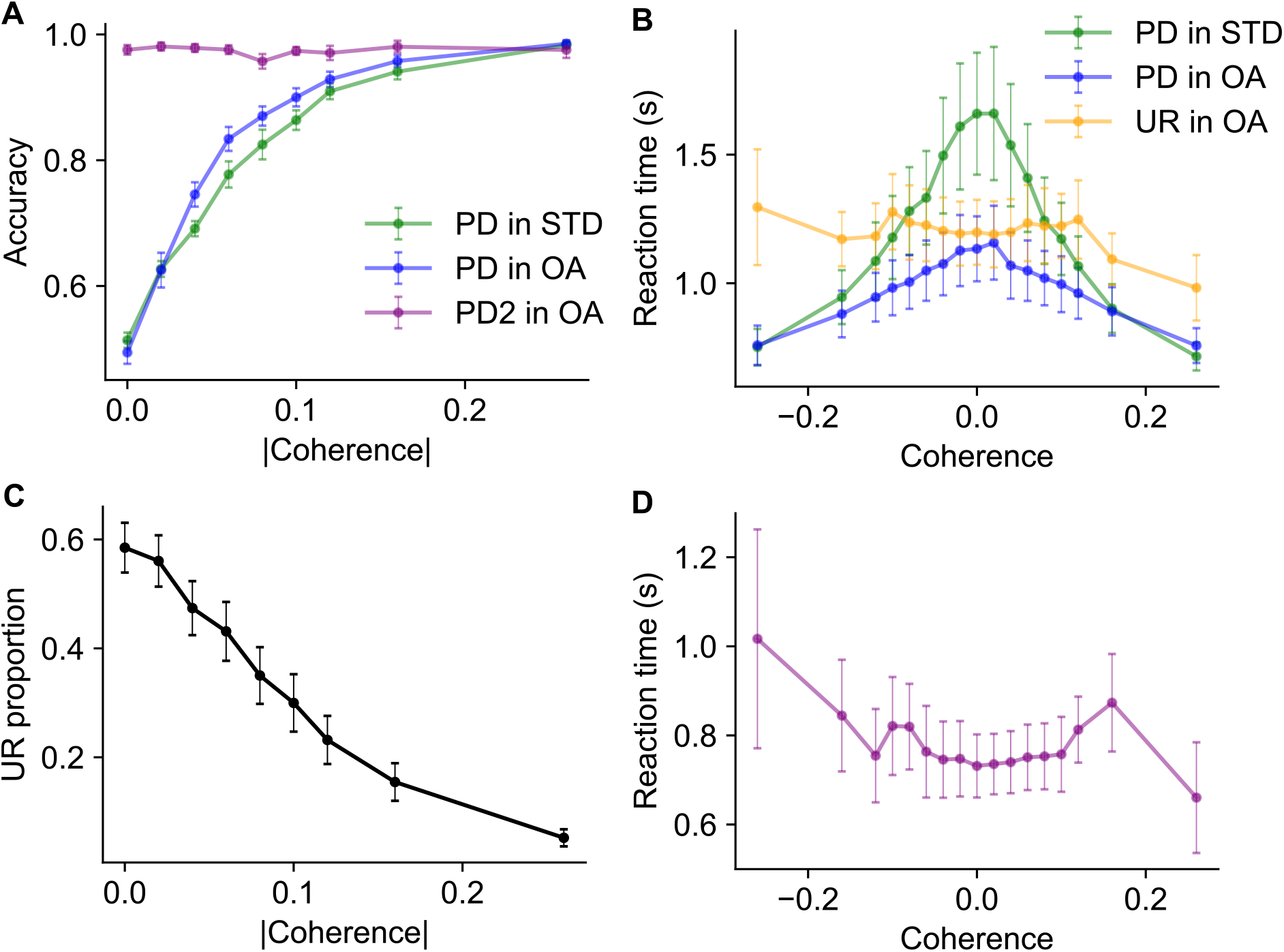
Subjects’ choice and reaction time patterns. **A)** Subjects’ accuracy plotted against the stimulus motion coherence for the perceptual decisions (PD) in the standard task (green), the direct PDs when the assistance option was available but not chosen (blue), and the perceptual decisions (PD2) after making the UR and viewing the second stimulus (yellow). **B)** The reaction times of PD (blue and green) and UR (yellow). **C)** The proportion of UR plotted against the first stimulus’s motion coherence. **D**) The reaction times of PD2 plotted against the first stage motion coherence. Error bars are SEM across the subjects (n=10). STD: standard task, OA: optional assistance task.

The population average pattern may mask the variability among the subjects. In the **Supplementary Figures S1 and S2**, we plotted the accuracy, the reaction time, and the UR proportion for every subject that we tested. While their psychometric curves followed the same pattern as the subject average (**Supplementary Figure S1**, left column), there appeared to be significant variations in their reaction times and UR proportions (**Supplementary Figure S1**, center and right columns).

To illustrate the individual variability, two representative subjects are shown in **Figure 3**. Both subjects’ PD accuracies were comparable and were a function of the first stimulus’s strength (**Figure 3A,D**). Compared to the reaction times of the PDs in the standard task, the reaction times of the UR were largely similar in subject 1 (**Figure 3B**) but significantly longer in subject 2 (**Figure 3E**). Both subjects made fewer UR when motion coherence was high, although the proportion of UR was lower in subject 2 (**Figure 3C,F**).

**Figure 3.**
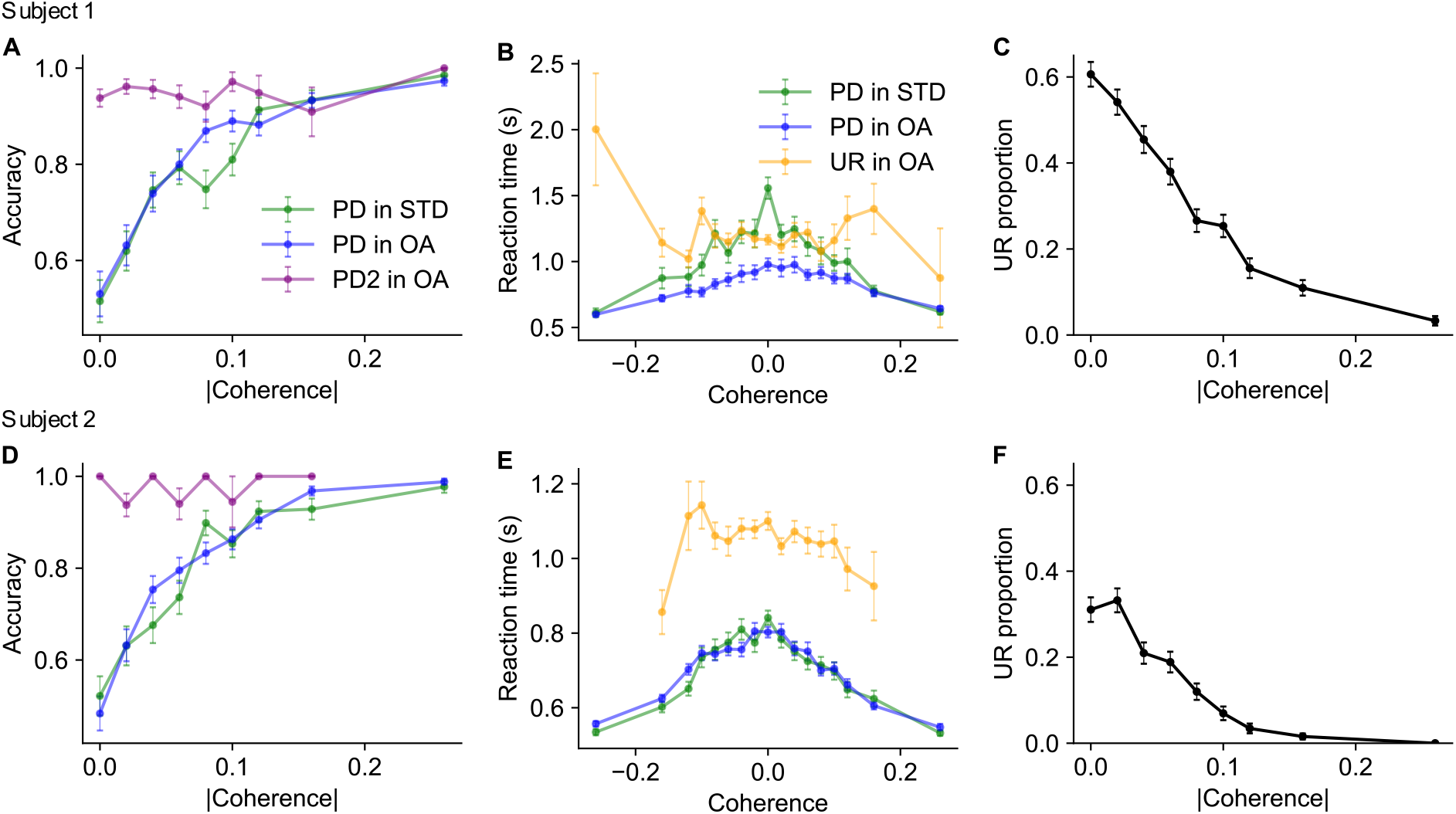
Performance and reaction times of two representative subjects. **A, B, C**) Representative subject 1’s accuracy, reaction time, and UR proportion plotted against motion coherence. Green: PD in the standard task, blue: PD in the optional assistance task, purple: PD2 in the optional assistance task. **D, E, F**) Representative subject 2. Green: PD in the standard task, blue: PD in the optional assistance task, yellow: UR in the optional assistance task. Error bars are SEM across trials in **A, B, D, E**.

### Model

To explain the findings from the behavioral experiment, we created a model inspired by an optimal policy decision model (OPDM) (Tajima et al., 2016, 2019). In the model, at each moment when performing the optional assistance task, the subject evaluates the expected returns from four alternatives, which are waiting for further evidence, choosing the left option, choosing the right option, and choosing the assistance option. The expected returns are computed based on the accumulated evidence, and the subject makes her choice to maximize her return.

We model the accumulation of evidence as an unbounded drift-diffusion process. The momentary evidence has a normal distribution 𝒩(*kzδt, σ*^2^*δt*), where *δt* is the size of each time step, *k* is the drift rate, and *z* is the motion coherence. The noise in the momentary evidence is independent across time as in the drift-diffusion model. The model assumes that the decision-maker holds a categorically distributed prior belief on the coherence strength *z*∼ Cat(***ξ, p***), where ***p*** is the probability of seeing coherence ***ξ*** in the current trial. The probabilities of seeing different motion coherences are uniform. Under these assumptions, after observing the momentary evidence over time, the subject updates her posterior belief about the true coherence level *z*.

The value function of the first stage *V*_1_(*t*_1_, *x*_1_) at time *t*_1_ when the subject has accumulated evidence *x*_1_ can be described with a Bellman equation:

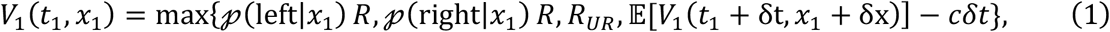

where *R* is the reward for a correct left-right response without seeking the assistance, 𝓅(left|*x*_1_) and 𝓅(right|*x*_1_) are the probabilities of the left and right choice being correct given evidence *x*_1_, *R*_*UR*_ is the reward for a correct response after seeking assistance (see Methods), and *c* is the time cost for waiting. We assume *R*_*UR*_ = 𝔼*V*_2_(0,0) − *f*, where 𝔼*V*_2_(0,0) is the second stage expected value, and *f* is a constant that reflects the cost of switching to the second stimulus. We also assume that *V*_2_(*t, x*) is calculated using the same drift-diffusion model parameters as in the first stage and the evidence accumulated during the first stage is not carried over. The subject’s goal is to choose the action that maximizes the value function *V*_1_(*t*_1_, *x*_1_). The perceptual decisions are based on the posterior belief on *z*. As the evidence is accumulated, the posterior belief on *z* evolves.

Based on the value functions, we plotted the optimal policy in the time-evidence plane (*t, x*) (**Figure 4A**) and the corresponding value function in **Figure 4B**. A trial starts at the center left of the time-evidence plane where the evidence and the time are 0. As evidence accumulates, the optimal policy drifts from the gray region of waiting to either the left decision region (above the upper boundary), the right decision region (below the lower boundary), or the UR region (wrapped inside the middle boundary). The UR region is enclosed inside the region for waiting but grows larger as time evolves.

**Figure 4.**
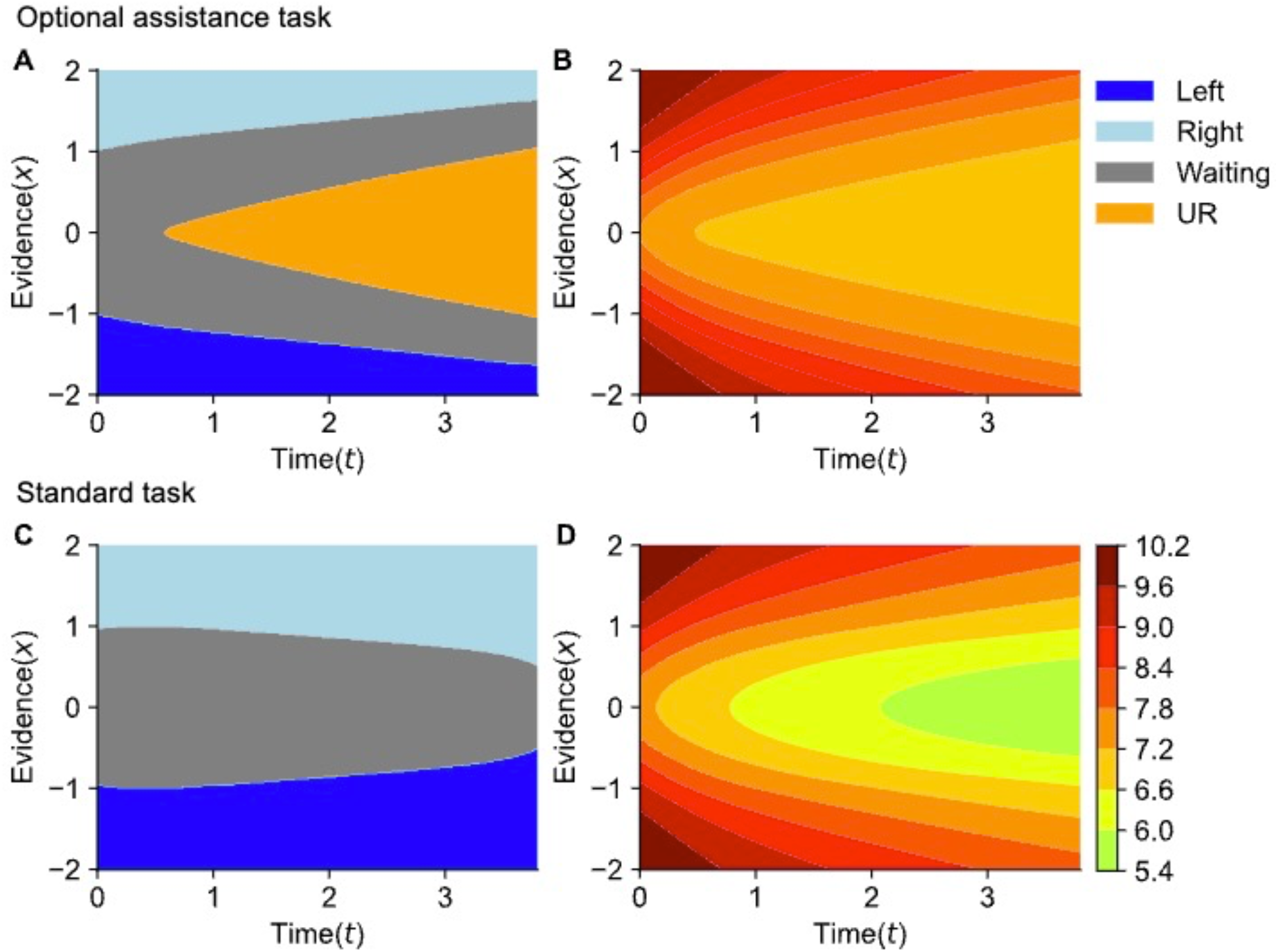
Optimal policy and its value function. **A**) Optimal policy plotted in the time-evidence plane for the optional assistance task. The color at each location indicates the optimal policy. **B**) The value function plotted in the time-evidence plane for the optional assistance task. The color depicts the value. **C**) Optimal policy plotted in the time-evidence plane for the standard task. **D**) The value function plotted in the time-evidence plane for the standard task. The default parameter setting is used unless otherwise mentioned.

For the standard task, in which the assistance option is not available, the value function reduces to:

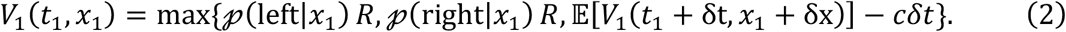

We plotted the optimal policy and the value function similarly for the standard task (**Figure 4C, D**). The optimal policy for the standard task only includes the left choice, the right choice, and the waiting. Therefore, one sees only a gray waiting region sandwiched between the upper and lower boundaries, corresponding to the left and right choices, respectively (**Figure 4C**). A low-value area appears in the right part of the value function’s heat map’s center region (**Figure 4D**, green). This is when the subjects have to make decisions based on only weak evidence.

With different parameter settings, the model replicated different subjects’ choice and reaction-time patterns. In particular, with two different settings of switching costs for UR *f*, **Figure 5** showed two simulations that produced choice and reaction time patterns similar to the representative subjects in **Figure 3**. In both settings, the model made more accurate PDs in the optional assistance task than in the standard task (**Figure 5A, D**). This difference was subtle at the level of individuals in **Figure 3**, presumably due to the small numbers of trials, but significant in the subject average (**Figure 2**). In addition, the difference was slightly more substantial in the first setting. When we used the default parameter setting (see Methods), the reaction times of the UR were comparable to those of the PD in the standard task (**Figure 5B**). This is comparable to representative subject 1 (**Figure 3B**).

**Figure 5.**
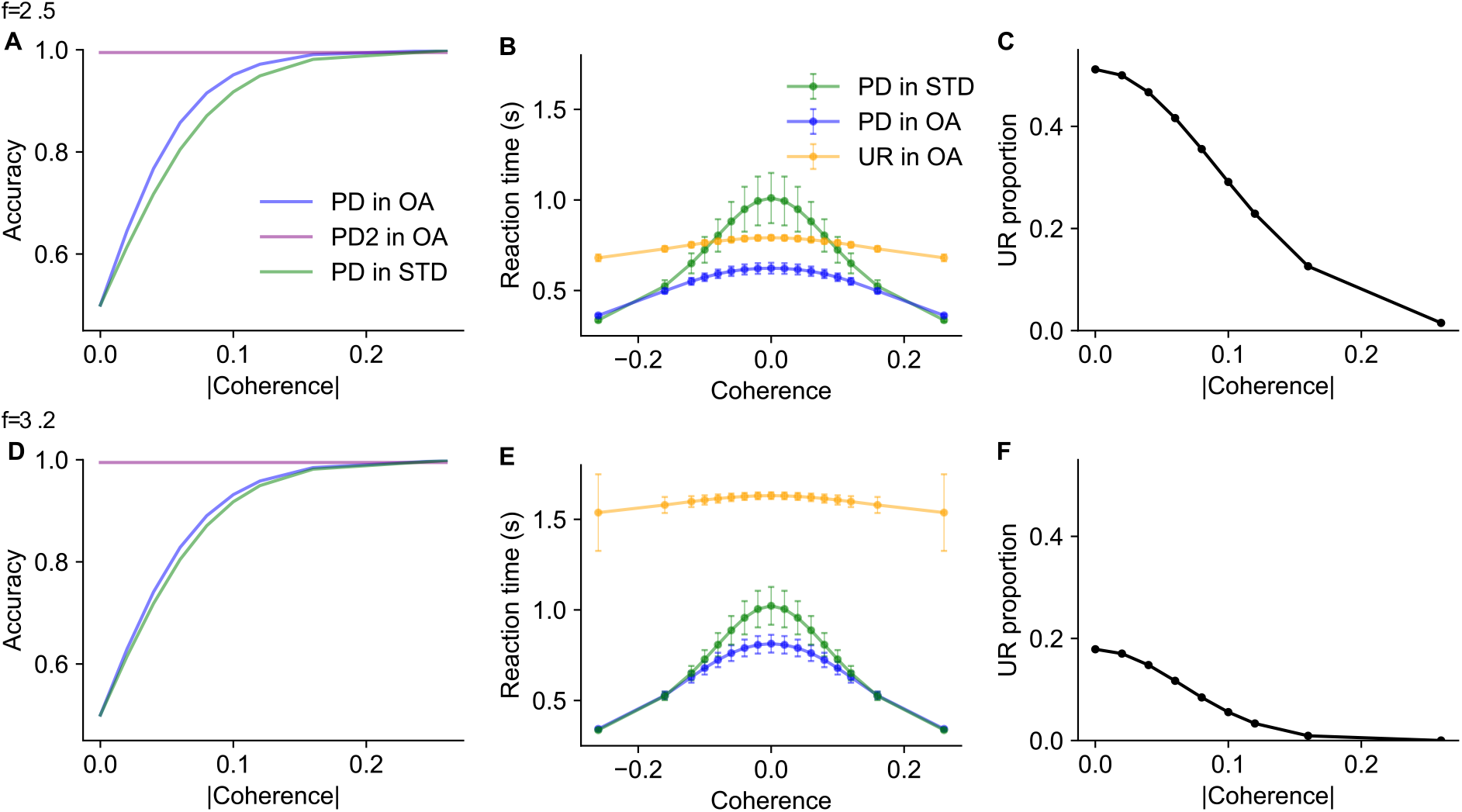
Model simulations. **A**) Accuracy, **B**) Reaction time, and **C**) UR proportion in the model with default parameter settings (*f* = 2.5). **D**) Accuracy, **E**) Reaction time and **F**) uncertainty response proportion (*f* = 3.2). PD: perceptual decision, PD2: perceptual decision in the second stage, UR: uncertainty response, STD: standard task, OA: optional assistance task.

When we raised *f* to 3.2 (see Methods), the model produced a reaction time pattern similar to representative subject 2’s (**Figure 3E, 5E**). The reaction times for the UR were substantially longer than for the reaction times for the PD in both tasks. A larger switching cost for UR not only makes choosing optional assistance less attractive but also causes the waiting to be more cost-effective. Hence, the uncertainty responses are slower. In addition, the proportions of UR are also lower with a higher *f* (**Figure 5C,F**).

Notably, although the UR’s reaction time curves produced by the model (**Figure 5B,E**, yellow) were flatter than the PD’s reaction time curves (**Figure 5B,E**, blue and green), they both concaved downwards. Reaction times were slightly faster for higher motion coherences in the model. Yet, the difference was small and become insignificant when tested with a moderate number of trials. We ran model simulations with 2,500 trials, a number comparable to what was used in the behavioral experiment (**Supplementary Table S1**), and varied the model parameters across a wide range. We found the correlations between the UR reaction times and motion coherences were much less likely to be significant (**Supplementary Figure S3**, yellow trace), while the reaction times for the PD almost always correlated with the motion coherences (**Supplementary Figure S3**, blue trace).

Comparing the two representative subjects in **Figure 3** and the model simulation results in **Figure 5** suggested that the accuracies, reaction times, and UR proportions may be correlated across the subjects, and the varying model parameters could account for the individual differences. This is indeed true. We first calculated the difference between the accuracies of the PD in the optional assistance task and the PD in the standard task for each subject. They correlated with the proportions of UR (Spearman *r* = 0.85, *p* < 0.01, see **Figure 6A**). Moreover, the subjects’ overall reaction time difference between their PD and their UR in the optional assistance task also correlated with the UR proportions negatively with an upward concavity (Spearman *r* = 0.92, *p* < 0.01, see **Figure 6B**). Subjects who made a larger number of URs also exhibited a greater improvement of PD accuracy and faster UR responses in the optional assistance task. Both patterns can be captured by adjusting *f*, the switching cost for UR, in our model (**Figure 6C,D**). In addition, similar patterns can also be reproduced by varying *R*, the reward for correct PDs (**Supplementary Figure S4A,B**). On the other hand, varying the cost for waiting *c* or the drift rate *k* produced slightly different patterns of the correlations between the UR proportion and the improvements in the PD accuracy, suggesting each alone was not sufficient to account for our data (**Supplementary Figure S4C,D,E,F**).

**Figure 6.**
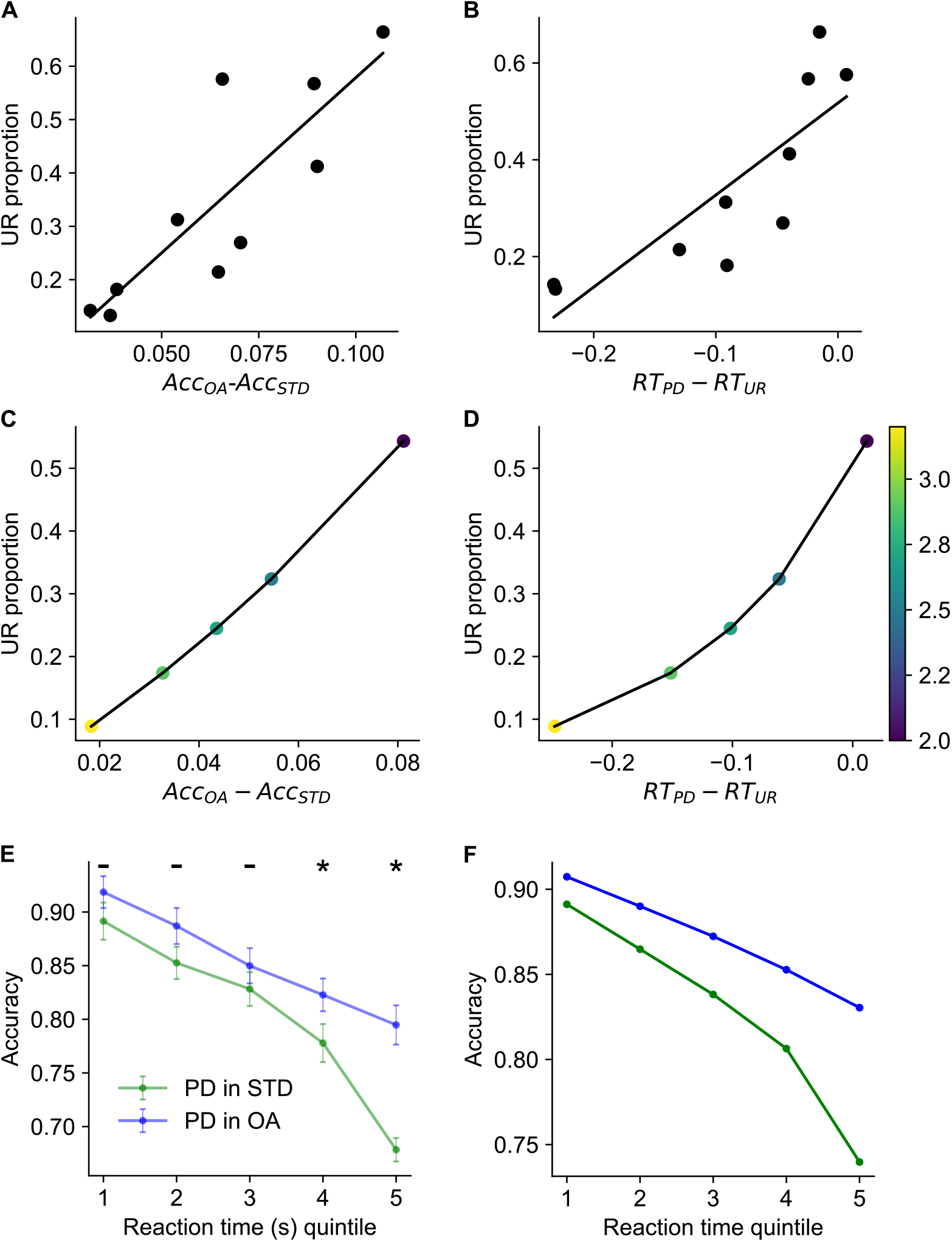
Interactions among UR proportion, reaction time, and accuracy. **A)** The improvement of the perceptual decision accuracy in the optional assistance task correlated with the UR proportion. Each data point is a subject. The line is a linear fit. **B)** The difference between the PD and the UR’s reaction time was correlated with the UR proportion in the optional assistance task. Each data point is a subject. The line is a linear fit. **C**,**D)** With different costs *f* for UR, indicated by the color from the scale bar on the right, the model reproduces the pattern in **A** and **B. E)** PD accuracy in the standard and the optional assistance tasks, plotted against the reaction time. To average across the subjects, we divided the reaction times of each subject into quintiles. Error bars indicate SEM across subjects. *p<0.05, Wilcoxon test. **F)** Model simulation for **E**. PD: perceptual decision, UR: uncertainty response, STD: standard task, OA: optional assistance task.

The presence of the UR region in the evidence-time plane (**Figure 4A**) leads to another interesting observation. The evidence required for the perceptual decisions increases over time as the UR region expands. This means that more evidence is required for perceptual decisions with long reaction times, and the typical negative correlation between PD’s accuracy and reaction time observed in the standard task may be alleviated by the presence of UR in the optional assistance task. To verify this prediction, we plotted the subjects’ PD accuracy against their reaction time separately in the two tasks (**Figure 6E, Supplementary Figure S5**). The decrease of accuracy was smaller in the optional assistance task. The performance differences did not reach statistical significance in the first three reaction time quintiles. This is easy to understand with the model, as the UR region does not exist when the reaction time is short (**Figure 4A**). Model simulations well captured this trend (**Figure 6F**).

## Discussion

The notion of (un)certainty and confidence is closely related. Here, we follow the nomenclature proposed by (Pouget et al., 2016). They defined confidence as the probability that a decision is correct given the evidence. As the quantity studied here is not based on a decision, we chose to use the term *uncertainty* to emphasize this aspect.

Our study uses a reaction time task to measure uncertainty and to ensure the uncertainty estimation is not based on decisions. Previous studies either required the subjects to indicate their decisions in addition to the confidence (Kiani et al., 2014; Lak et al., 2014) or probed the confidence only after the presentation of sensory stimuli at fixed durations, thus allowing the subjects to form covert decisions (Kiani & Shadlen, 2009; Komura et al., 2013). Only with a reaction-time design, subjects may form uncertainty decisions before perceptual decisions and the two decision processes can be distinguished.

We observed clear evidence that the uncertainty estimation process occurred in parallel with the perceptual decision process. If the uncertainty estimations are made after and based on covert PDs, we would expect the reaction times of the UR to follow a dependency on the motion coherence as the PD reaction times do: fast for high coherence motion stimuli and slow for low coherence motion stimuli. This is not consistent with our data. The reaction times for the UR did not significantly correlate with the motion coherence. This result is not due to the subjects’ failure of making appropriate uncertainty estimations. The proportions of UR strongly correlated with the motion coherence, reflecting that the degree of uncertainty increased as the motion coherence grew weak. This is consistent with many previous studies (Kiani & Shadlen, 2009; Komura et al., 2013; Shields et al., 1997; D. J. Smith, 2009), although they did not measure the reaction time of confidence or (un)certainty along. The reaction time pattern of the uncertainty responses in our results also excludes the proposal that uncertainty estimation may be based on the elapsed time during decision making (Kiani et al., 2014).

It is noteworthy that our model predicts a weak correlation between the reaction time of uncertainty response and the motion coherence. However, the predicted correlation is much weaker than that between the PD reaction time and the motion coherence, and cannot be detected statistically when there is significant behavior noise and the number of trials is small (**Supplementary Figures S2 and S3**), as in the case of the actual experiments. Nevertheless, our results do not exclude an alternative scenario in which the URs were based on an estimation of motion coherence against a reference point. This would lead to shorter reaction times for both the low and the high extremes of motion coherences but longer reaction times for coherences near the reference point. Future experiments with a more extensive data set may be used to test this hypothesis.

The current model assumes that both the uncertainty estimation and the perceptual decision are based on the same evidence accumulation process, therefore the same neural circuitry may be underlying both computations. Consistent with this idea, it has been shown that neuronal activity in the lateral intraparietal area (LIP), which encode evidence accumulation signal during decision making, reflected confidence (Kiani & Shadlen, 2009). More recent studies using microstimulation or optogenetics tools to manipulate neuronal activities in the sensory area also led to changes in confidence, providing further evidence (Fetsch et al., 2014, 2018). It has also been shown that a biologically realistic neural network may compute both perceptual decision and confidence estimation simultaneously, and its neurons demonstrated response patterns similar to those of LIP neurons (Wei & Wang, 2015).

Our model makes its choices by comparing the values of different options, and it would be interesting to ponder how the brain might implement the value computation based on evidence accumulation. Ramping activities that reflect evidence accumulation have been reported in many brain regions, including the prefrontal cortex (Kim & Shadlen, 1999), the posterior parietal cortex (Roitman & Shadlen, 2002; Yang & Shadlen, 2007), the basal ganglia (Ding & Gold, 2010; Doi et al., 2020), the frontal eye field (Gold & Shadlen, 2000) and the superior colliculus (Jun et al., 2021) during decision making. In addition, neurons in the orbitofrontal cortex and the dorsolateral prefrontal cortex have been shown to encode values associated with sensory stimuli and accumulated evidence during decision making (Lin et al., 2020). Therefore, it is reasonable to postulate that the structures in the reward circuitry, areas in the prefrontal cortex in particular, may be involved in computing values associated with accumulated evidence and uncertainty responses, by receiving the inputs from the brain regions, such as LIP, where evidence is accumulated, and thereby contribute to concurrent uncertainty estimation. Consistent with this idea, recent rodent studies pointed to the orbitofrontal cortex as a key player in confidence estimation (Kepecs et al., 2008; Lak et al., 2014; Masset et al., 2020).

Finally, our model suggests that the uncertainty estimation can be resolved at a perceptual level. A recent study introduced a value-based Bayesian framework based on partially observable Markov decision processes to explain the results in the earlier confidence study (Khalvati et al., 2021). In the framework, both the perceptual decisions and the opt-out decisions were based on the same hidden belief updated with the sensory inputs. Therefore, the framework provided a different approach to derive confidence, again, at a perceptual level. These studies question whether confidence and (un)certainty estimation studied by many can be regarded as meta-cognition as claimed (Kiani et al., 2014; Kiani & Shadlen, 2009; Komura et al., 2013) and whether metacognition, in this case, is useful to describe the computation of confidence or (un)certainty in the brain during decision making.

## Data and code availability

Data and code are available from the corresponding author on reasonable request.

## Acknowledgements

We thank Roozbeh Kiani for discussions and comments during the study, and Ninglong Xu and Ningyu Zhang for their comments during the preparation of the manuscript. We also thank Zhewei Zhang, Zhongqiao Lin, Chechang Nie, Yang Xie, Wei Kong, Lu Yu for their help in all phases of the study. The authors declare no competing financial or nonfinancial interests.

## Author contributions

T.Y. designed the study. X.L., R.S. and Y.C. performed the experiments and collected the data. X.L. and R.S. analysed the data. X.L. designed the model. X.L. and T.Y. wrote the paper.

## Methods

### Subjects

We collected behavior data from 10 human subjects (18–28 yr of age; 6 males) who had normal or corrected-to-normal vision. Two subjects were the authors. The others were graduate or college students. All experimental procedures followed the protocol approved by the Ethics Committee of Shanghai Institutes for Biological Sciences, Chinese Academy of Sciences (Shanghai, China).

### Behavior Task

The subjects were seated in a quiet dark room with their head stabilized by a chin rest. A 17” LCD monitor (refresh rate: 60 Hz; screen resolution: 1440*900) was located 51 cm in front of the subjects, spanning 41cm horizontal and 24cm vertical of their central visual field. Subjects interacted with the program by pressing arrow keys on a keyboard. The behavioral tasks, stimulus presentation, and collection of behavioral data were implemented with PsychoPy3 (2020.1.3). Data were analyzed offline with Python3.

The standard task was a reaction-time version of the classic random dot motion discrimination task. At the beginning of each trial, two white arrows appeared at the left and right peripheral regions on the monitor. Subjects needed to press and hold down the down arrow key to start a trial and view the random-dot stimuli. The random dot motion patterns were created using PsychoPy3’s built-in functions, displayed in a virtual aperture (7.5cm diameter) at the center of the screen. Subjects were asked to judge whether, on average, the motion direction of dots was toward left or right. The stimulus lasted at most for 10 seconds, but the subjects were allowed to indicate their decisions whenever ready by releasing the down arrow key and pressing the left or right arrow key within 200 ms. Afterward, a green checkmark or a red cross would appear on the corresponding arrow’s position to indicate a correct or an erroneous response. The reaction time was defined as the duration between the motion stimulus onset and the down arrow key release.

Motion strengths and directions of dots varied randomly from trial to trial. The motion strength of dots in each trial was selected randomly from a coherence gradient (0%, 2%, 4%, 6%, 8%, 10%, 12%, 16%, 26%) with equal probabilities. Motion directions were either toward the left or the right at equal probabilities. At 0% coherence, the direction was artificially defined, and all dots moved in random directions.

In the optional assistance task, in addition to the two arrows on the left and right, a white downward arrow appeared at the bottom region of the monitor. The subject instruction (originally in Chinese) was: “If you are confident enough to make a correct perceptual decision, you can press the left or the right key whenever you are ready. If you are not sure about the answer, you can release the down key and quickly press and hold it again to view an easier stimulus and make a choice accordingly afterward.” The second stimulus’s motion strength was fixed at 26%, and its direction was always consistent with the first stimulus.

After viewing the second stimulus, the subject could release the down key and quickly press the left or the right key to report their decision. The second stimulus lasted also maximally 10 seconds. The reaction times of the first stage responses are defined as the duration between the onset of the first stimulus and the release of the down arrow key. The reaction times of the second stage decisions are defined as the duration between the onset of the second stimulus and the second release of the down arrow key.

All subjects had experience in various versions of random dots motion direction discrimination tasks. They went through at least 4 days of training in our task before data collection. The data were then collected in the next 5 days. The trials were separated into alternating blocks of the standard task (block size =50) and the optional assistance task (block size =100). In total, 2500 optional assistance task trials and 1250 standard task trials were collected for each subject. Missed responses and responses that were made too fast (<250ms) were excluded from the analyses. The actual numbers of trials used in the analyses for each subject were indicated in **Supplementary Table S1**.

### The model

#### Structure of evidence and evidence accumulation

Our model is adapted from an optimal policy decision model (Tajima et al., 2016, 2019). We define a categorical distribution to indicate different coherence levels *z*. Negative coherences indicate leftward motions, and positive coherences indicate rightward motions. The subjects are assumed to know the distribution of *z*, but not the exact coherence level for each trial. The coherence distribution is given by *z* ∈ Cat (***ξ, p***), where ***ξ*** = {*ξ*^(1)^, …, *ξ*^(*M*)^} is the set of coherences, ***p*** = {*p*^(1)^, …, *p*^(*M*)^} is the set of the probabilities of being shown, and Cat stands for the categorical distribution. Thus, *z*_1_∼Cat (***ξ***_1_, ***p***_1_) and *z*_2_∼Cat (***ξ***_2_, ***p***_2_) are the first and second stage coherence distributions respectively.

For the standard task and the first stage of the optional assistance task, the coherence levels and their probabilities are

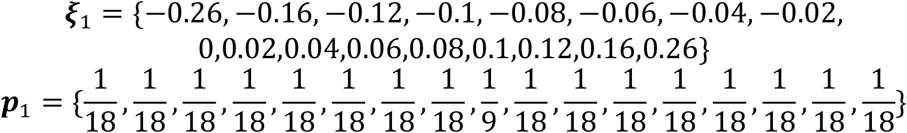

For the second stage of the optional assistance task, ***ξ***_2_ = {−0.26,0.26} and 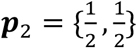.

We assume that the subject observes noisy momentary evidence *δx*^(*i*)^ with a normal distribution 𝒩(*kzδt, σ*^2^*δt*) in time-step *i* of duration *δt*. The subject accumulates the evidence 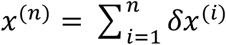 up to time *t*^(*n*)^ = *nδt*, and the posterior probability of *z* is

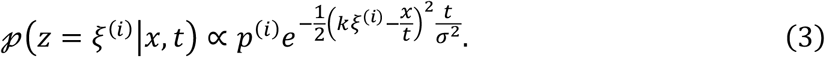

#### Accuracy for direction discrimination

The subject’s posterior belief about the accuracy of choosing the left direction in the first stage is

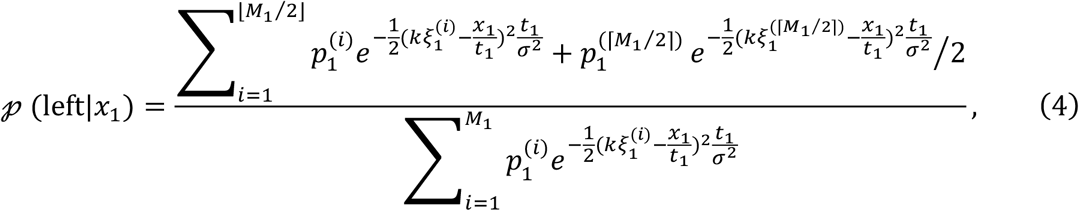

and the posterior belief about the accuracy of choosing the right direction is

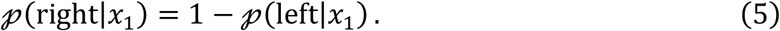

The accuracies of left or right choices at zero-coherence are equally 0.5.

In the second stage, the subject’s posterior belief about the accuracy of choosing the left direction can be given by

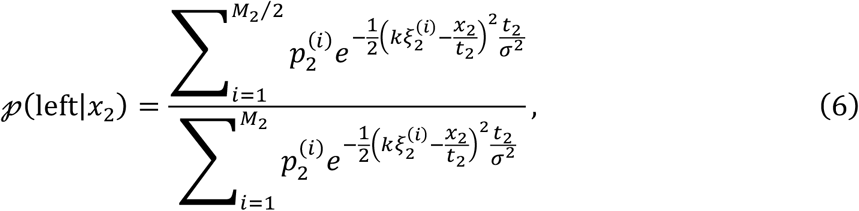

and

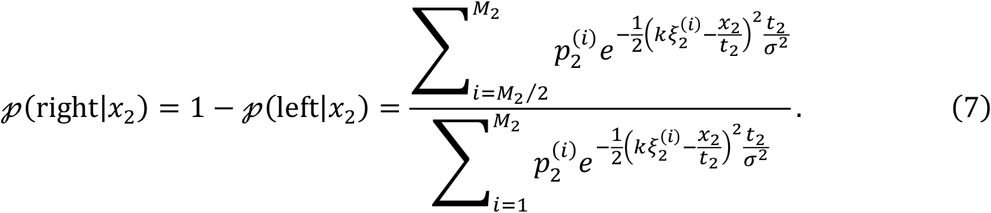

#### Bellman equation

We use the Bellman equations to form subjects’ decision-making boundaries. The value functions can be described as

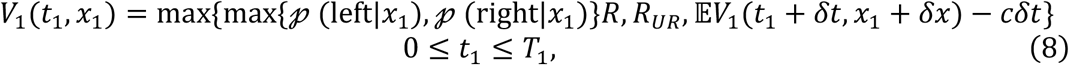

for the first stage, and

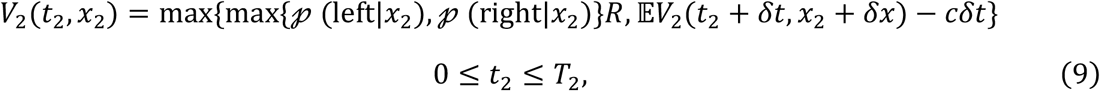

for the second stage. Here, *R* is the reward for a correct perceptual decision, *c* is the cost per time step. 𝔼*V*_1_(*t, x*) and 𝔼*V*_2_(*t, x*) are the expected value at time *t* with accumulated evidence *x. R*_*UR*_ is the total value for switching to the second stage: *R*_*UR*_ = 𝔼*V*_2_(0,0) − *f*, where 𝔼*V*_2_(0,0) is the second stage expected value and *f* is the switching cost. We solve the Bellman equation using the grid method and backward induction. The grid range is [−*g, g*] for *x* and [0, *T*] for *t*, and *g* is set to 𝓅(right|*T, g*) = 0.2. The expected value functions 𝔼*V*_1_(*t, x*) and 𝔼*V*_2_(*t, x*) are calculated as follows,

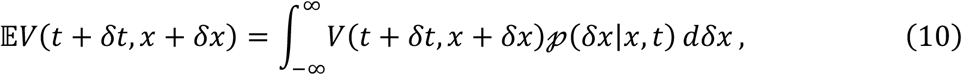

where 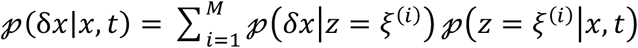. The integration is calculated numerically with *δx* ∈ [*x* − 3*σ, x* + 3*σ*] in grids, and *δt* is set to 0.01.

#### Decision Boundaries and Markov chain approximations

The optimal policy of the Bellman equations identifies the decision boundaries choosing left, right, uncertainty response, or waiting in the evidence plane (*t,x*). The evidence structure given above is the same as the drift-diffusion model. Hence the first passage time on these boundaries can be computed in the same way as the drift-diffusion model. We adopt a simple matrix method (Diederich & Busemeyer, 2003; Ditterich, 2006) to approximate the reaction time distribution instead of directly simulating trials for more accurate and time-saving results. We extend their methods by adding an UR decision boundary embedded inside the waiting region, and there are three absorbing states, which correspond to the left, the right, and the UR decisions.

In the Markov approximation, we discretize time *t* and accumulated evidence *x* in the same way as the grid method in the Bellman equation. Hence, the decision boundaries formed with the Bellman equation can be directly taken into the Markov chain method. The starting point of the drift-diffusion is zero. At each time step, the transition matrix is derived as follows.

If the state is inside the UR region, i.e., 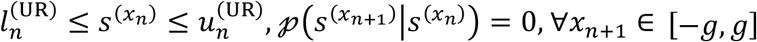 and 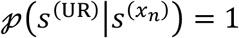, where 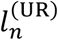 and 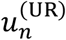 are the Markov states that correspond to the lower and upper branches of the uncertainty response boundary. *s*^(UR)^ is the UR absorbing state.

Let 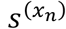 be the grid state that is closest to the continuously accumulated evidence *x*_*n*_. Note that the states are integers between 0 and *N*_*s*_ = 300 and share the same grid range and numbers described in the section above. Let *y*_*i*_ = *Φ*(*h*_*i*_) − *Φ*(*h*_*i*+1_), where *Φ* is cumlative probability of the standard normal distribution, *h*_(_ is the *i-*th component of the even divisions of 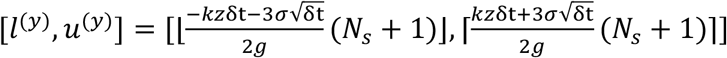, and the size of each division is 1. Let *l*_*n*_ and *u*_*n*_ stands for the states of the lower and the upper branches of the UR boundary at time *n* respectively.

If the UR region exists at time step *n*, the transition probability from the current state 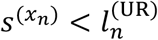 to the non-UR states can be calculated in four conditions: below the lower bound for the perceptual decision, at the lower bound, above or equal the lower branch of the UR boundary, or between the two bounds and in the waiting region:

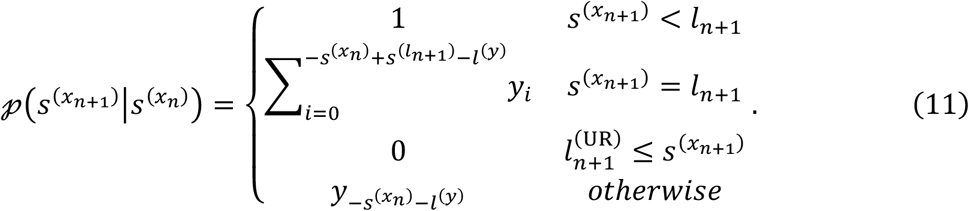

Similarly, the transition probability for 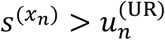 can be computed as:

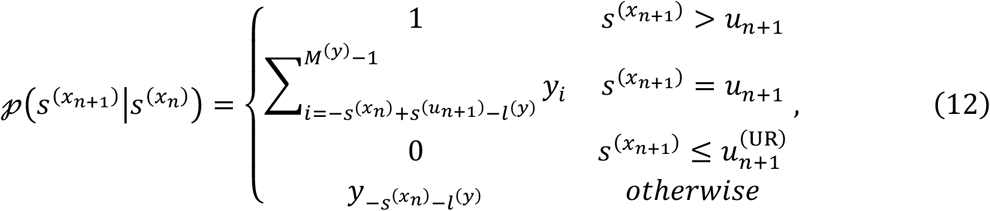

where *M*^(*y*)^ is the number of the divisions of [*l*^(*y*)^, *u*^(*y*)^].

When the current state is not in the UR region but may reach the UR region, the probability of transiting to the UR state is:

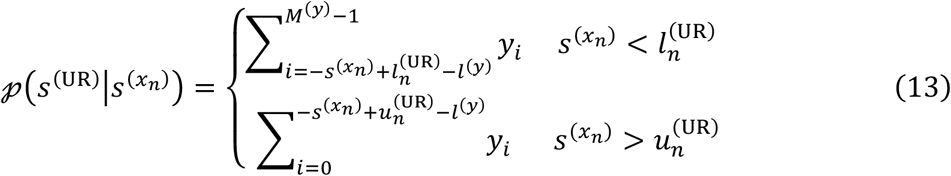

For the states inside UR region, 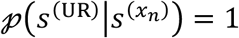. Also, 𝓅 (*s*^(*UR*)^|*s*^(*UR*)^) = 1.

If the state is not in the UR region at step *n*, but it might reach the UR region at the next time step, then

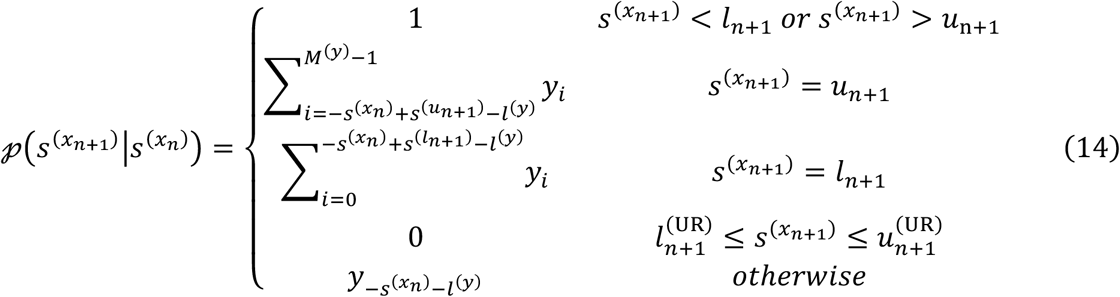

for 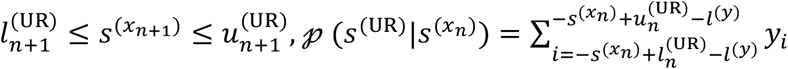 and for the UR state itself, 𝓅 (*s*^(UR)^|*s*^(UR)^) = 1.

By iterating each time point, we can get the distributions of reaction times for each of the choices 𝓅 (*t*, left|*z*), 𝓅 (*t*, right|*z*) and 𝓅 (*t*, UR|*z*).

#### Model parameter settings

The default parameter setting used in the model simulations in **Figures 4, 5, 6DEF, S3**, and **S4** is *k* = 12, *a* = 10, *c* = 1, *f* = 2.5, *σ* = 1, *T* = 4.

#### Simulation Analyses (Figures 5 and 6)

Using the Markov chain approximation procedure described above, we get three reaction time distributions for the left 𝓅 (*t*, left|*z*), the right 𝓅 (*t*, right|*z*), and the uncertainty responses 𝓅 (*t*, UR|*z*) at each coherence level *z*. In **Figure 5**, the accuracy is 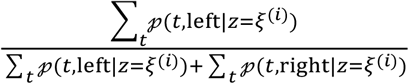, and the proportion of uncertainty responses is 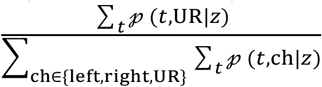, where *i* = {⌊*M* /2⌋, …, *M*}. The mean of the reaction times given a coherence level is RT(*z*) ≡ ∑_0_ *t* 𝓅(*t*|*z, choice*), and the variance is ∑_0_(*t* − RT (*z*))^2^𝓅 (*t*|*z, choice*), where choice ∈ {left, right, UR}.

In **Figure 6CD**, the overall proportion of uncertainty responses for each subject is 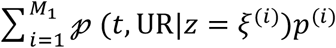, the accuracy is 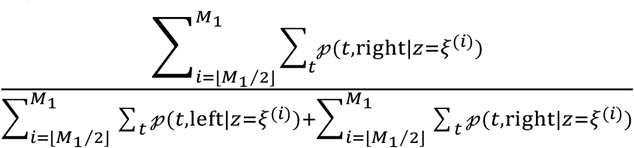 in **Figure 6C**, and the mean reaction time is 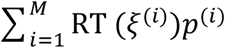 in **Figure 6D**.

To be compared across subjects, the perceptual decisions’ RTs across tasks for each subject are standardized in **Figure 6BE**. The trials from each subject are divided into quintiles based on the reaction time in the standard and the optional assistance tasks in **Figure 6E**. Accuracies in **Figure 6F** are similarly computer as in **Figure 6C**.

## Supplementary Table and Figures

**Table S1.** Trial numbers. Shown in the table are trial numbers for each task type and each subject used in the **Figures S1, S2, S3, S4, and S5** and their related analyses. Trial numbers for the direct left/right choices and the uncertainty responses under the uncertainty response task are separately listed.

| subjects | Standard Task | Optional Assistance Task |  |  | total |
| --- | --- | --- | --- | --- | --- |
|  |  | total | left/right | UR |  |
| **1** | 1201 | 2475 | 2025 | 450 | 3676 |
| **2** | 1216 | 2458 | 1931 | 527 | 3674 |
| **3** | 1234 | 2487 | 1710 | 777 | 3721 |
| **4** | 1234 | 2475 | 2123 | 352 | 3709 |
| **5** | 1238 | 2486 | 1461 | 1025 | 3724 |
| **6** | 1233 | 2494 | 1822 | 672 | 3727 |
| **7** | 1216 | 2459 | 2132 | 327 | 3675 |
| **8** | 1234 | 2474 | 831 | 1643 | 3708 |
| **9** | 1234 | 2466 | 1067 | 1399 | 3700 |
| **10** | 1236 | 2438 | 1034 | 1404 | 3674 |

**Figure S1.**
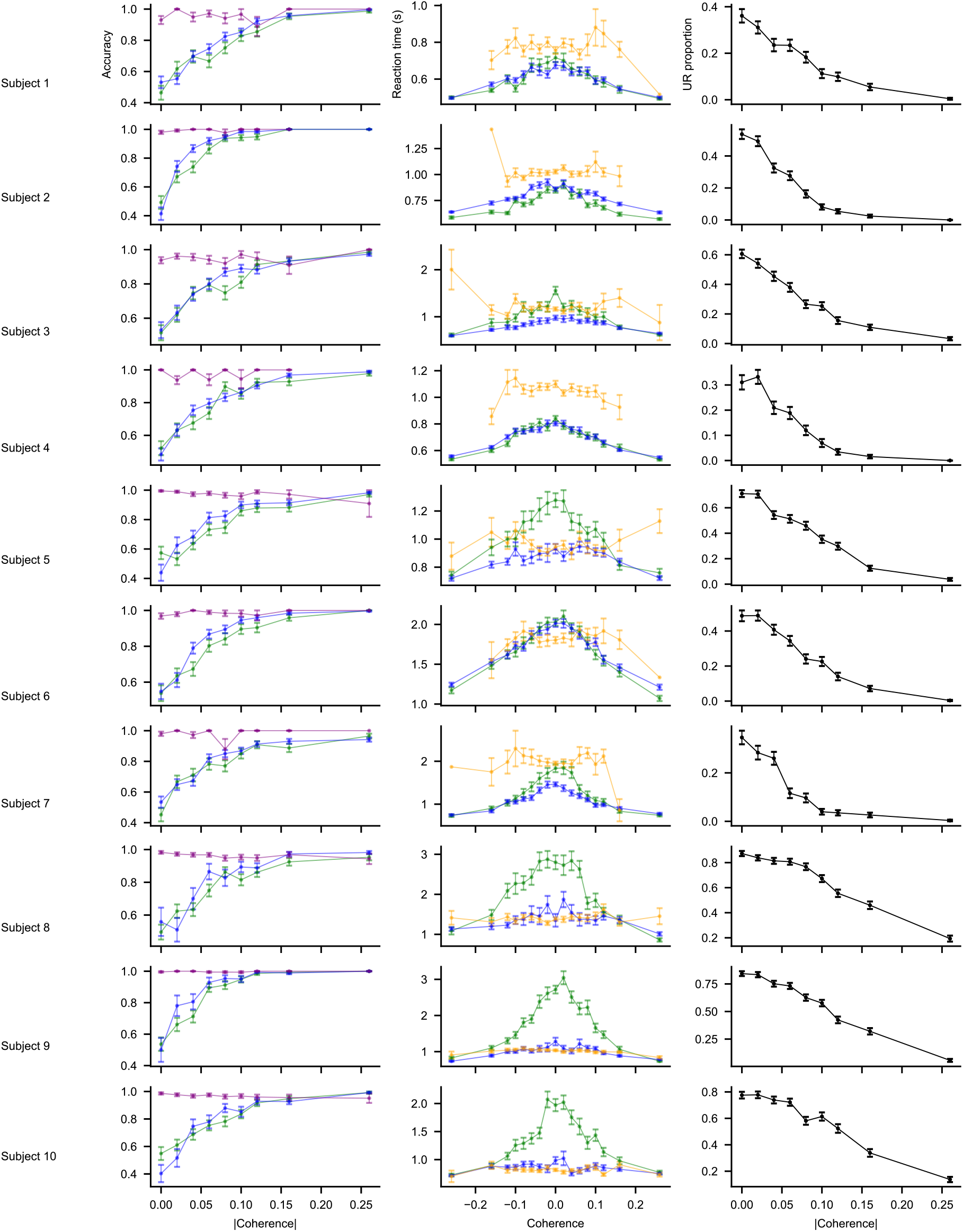
Each subject’s accuracies, reaction times, and proportions of UR plotted as a function of motion coherence. Left column: Accuracy. Green: standard task, blue: direct perceptual decisions in the optional assistance task, purple: perceptual decisions in the second stage in the optional assistance task. Center column: Reaction time. Green: perceptual decisions in the standard task, blue: direct perceptual decisions in the optional assistance task, orange: UR in the optional assistance task. Right column: Proportions of UR. Error bars indicate S.E.M. across trials. PD: perceptual decision, PD2: perceptual decision in the second stage, UR: uncertainty response, STD: standard task, OA: optional assistance task.

**Figure S2.**
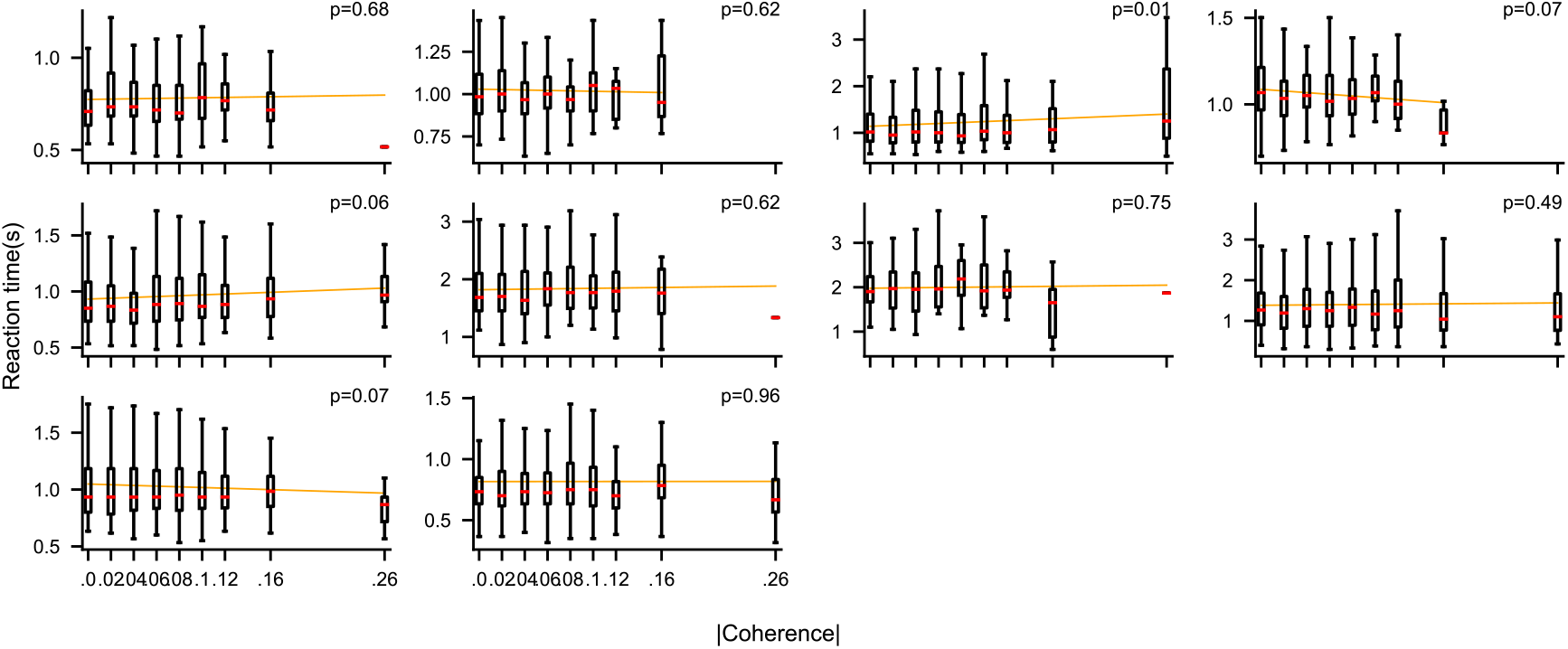
Reaction times of UR. The orange line is the reaction times regressed on coherence levels with the ordinary least square method. The boxes indicate the first and third quartiles of the reaction times for each coherence, and the upper and lower whiskers indicate the maximum and minimum reaction times. p values indicate the significance of the regressions.

**Figure S3.**
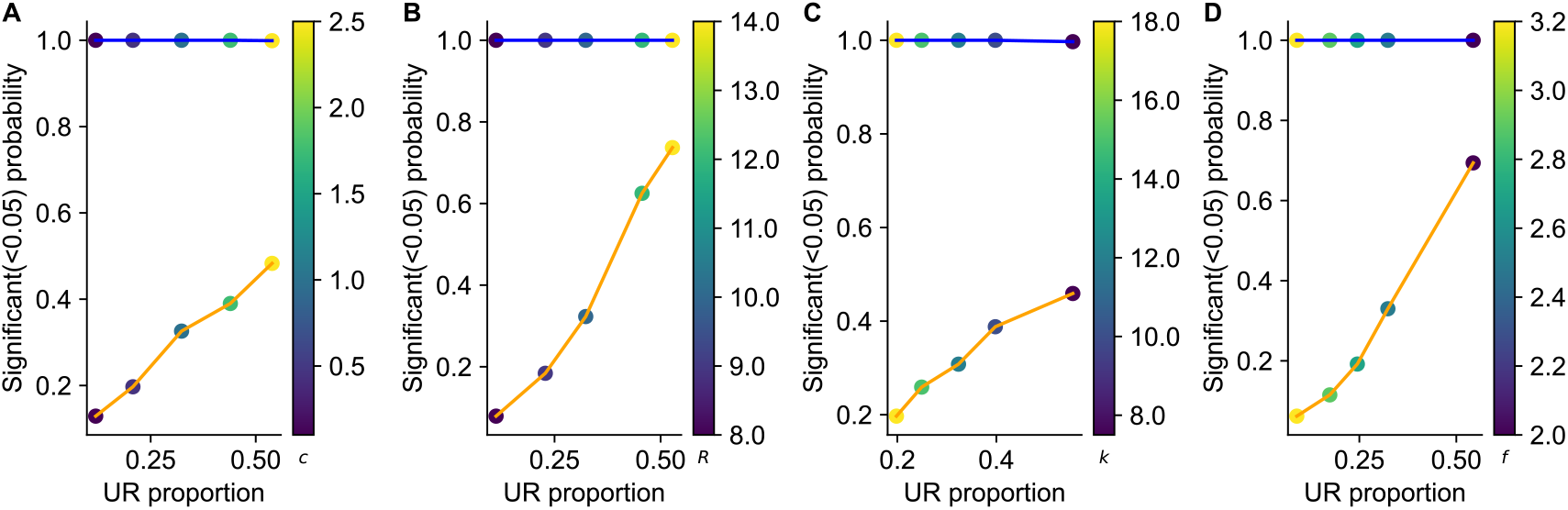
Proportions of simulations that produced significant correlations (p<0.05) between the motion coherence and the PD reaction time (blue) and between the motion coherence and the UR reaction time (yellow). In each panel, one of the model parameters was varied (*c, R, k, R*_*UR*_ in **A, B, C**, and **D**), and each data point corresponds to the average of 1000 simulations based on a parameter setting indicated by the color (2500 trials in each simulation).

**Figure S4.**
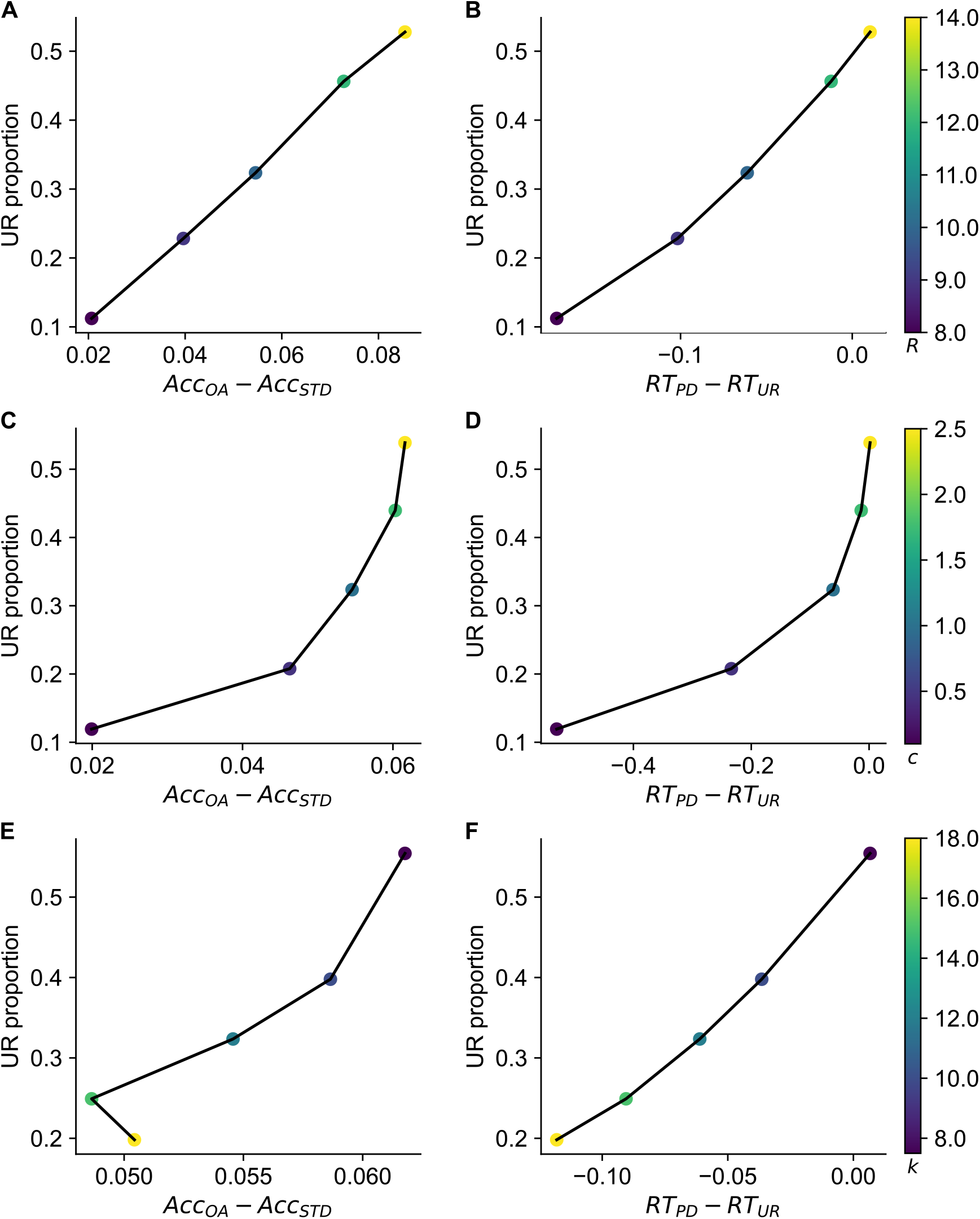
Interactions among UR proportion, reaction time, and accuracy. **A, B)** With different settings for the reward *R*, the model reproduces the patterns in **Figure 6A,B**, in which *f* is varied. **C**,**D)** Different settings for the cost for waiting *c*. The relationship between the UR proportion and the improvement of perceptual decision accuracy is much more curved. **E**,**F)** Different settings for the drift rate *k*. There is a non-monotonic relationship between the UR proportion and the improvement of perceptual decision accuracy. The colors of the dots and the vertical color scale bars indicate the parameter values.

**Figure S5.**
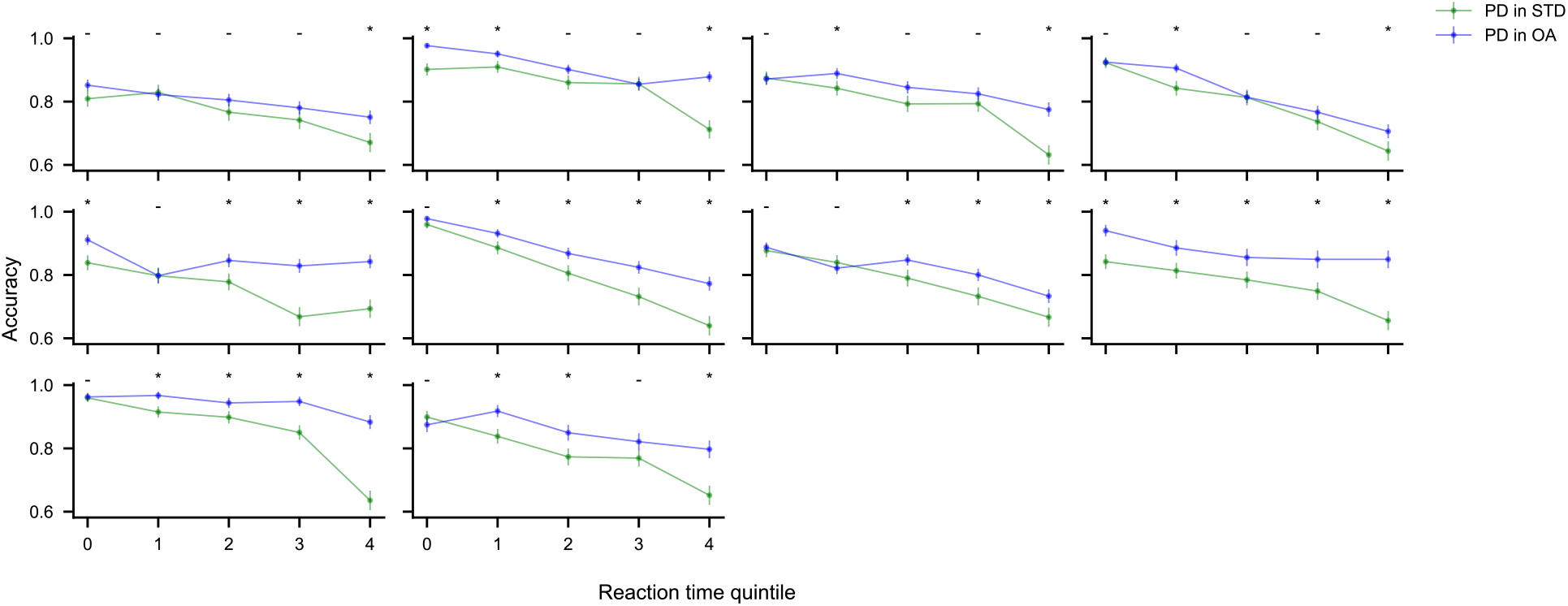
Accuracy plotted as a function of reaction time plotted for each subject. For each subject, the trials are grouped into quintiles by their reaction time. Error bars indicate S.E.M. Green: standard task, blue: optional assistance task. STD: standard task, OA: optional assistance task. * indicated p<0.05 (Wilcoxon test).

